# Disruption of *Brachypodium* Lichenase Alters Metabolism of Mixed-linkage Glucan and Starch

**DOI:** 10.1101/2021.08.20.457123

**Authors:** Mingzhu Fan, Jacob K. Jensen, Starla Zemelis-Durfee, Sang-Jin Kim, Jia-Yi Chan, Claudia M. Beaudry, Federica Brandizzi, Curtis G. Wilkerson

## Abstract

Mixed-linkage glucan (MLG), which is widely distributed in grasses, is a polysaccharide highly abundant in cell walls of grass endosperm and young vegetative tissues.
Lichenases are enzymes that hydrolyze MLG first identified in MLG-rich lichens. In this study, we identify a gene encoding a lichenase we name *Brachypodium distachyon LICHENASE 1* (*BdLCH1*), which is highly expressed in the endosperm of germinating seeds and coleoptiles and at lower amounts in mature shoots. RNA in situ hybridization showed that *BdLCH1* is primarily expressed in chlorenchyma cells of mature leaves and internodes. Disruption of BdLCH1 resulted in an eight-fold increase in MLG content in senesced leaves. Consistent with the in situ hybridization data, immunolocalization results showed that MLG was not removed in chlorenchyma cells of *lch1* mutants as it was in wild type and implicate the BdLCH1 enzyme in removing MLG in chlorenchyma cells in mature vegetative tissues. We also show that MLG accumulation in *lch1* mutants was resistant to dark induced degradation, and eight-week-old *lch1* plants showed a faster rate of starch breakdown than wild type in darkness. Our results suggest a role for BdLCH1 in modifying the cell wall to support highly metabolically active cells.

## INTRODUCTION

Walls of land plants contain cellulose, hemicellulose, pectin, lignin, and some structural proteins (Keegstra, 2010; Scheller and Ulvskov, 2010). MLG, which is a matrix polysaccharide that is widely distributed in commelinoid monocots, consists of unbranched and unsubstituted glucose residues and can be hydrolyzed into glucose by commercial hydrolases (Burton and Fincher, 2009). Thus, MLG is an easily fermentable cell wall polysaccharide and an ideal compound for the production of biofuels and glucose-derived bioproducts (Pauly and Keegstra, 2008). Plants with high amounts of MLG would therefore be valuable as industrial feedstocks. MLG is also likely involved in cell development. It has been reported that plant vegetative tissues accumulate MLG during cell expansion and a substantial amount of this MLG is subsequently removed from mature walls by MLG hydrolases (Kim et al., 2000; Carpita et al., 2001). So far, the mechanism of MLG removal during wall maturation is not clear nor is it known if all cell types participate in this process.

Overproduction of MLG synthases or disruption of MLG hydrolases are two likely strategies to increase the amount of MLG in plants. The cellulose synthase-like gene families *CSLF, CSLH, CSLJ* are involved in MLG biosynthesis with *CSLF6* encoding the primary synthase (Burton et al., 2006; Burton et al., 2008; Doblin et al., 2009; Little et al., 2018). Overexpression of *CSLF6* in barley and *B. distachyon* led to an increase in MLG, however the resulting plants exhibited detrimental effects on growth and development (Burton et al., 2011; Kim et al., 2018). Most of the barley *CSLF6* overexpression transformants could not survive until maturity. A similar phenotype was also observed when overexpressing *BdTHX1* in *B. distachyon*, a trihelix transcriptional factor that likely regulates *BdCSLF6* and a gene that encodes an MLG:xyloglucan endotransglucosylase BdXTH8 (Fan et al., 2018). The negative effect on plant growth is currently a barrier to increase MLG accumulation in the plant by increasing the rate of MLG production.

Another strategy to increase MLG content in grasses is disrupting endogenous MLG hydrolases, known as lichenases or (1,3;1,4)-β-d-glucan 4-glucanohydrolases (EC 3.2.1.73). These enzymes typically hydrolyze a β-(1,4)-d-glucosidic linkage that immediately follows a β-(1,3)-d-glucosidic linkage in MLG (McCleary, 1988). In grasses, two lichenase isoenzymes, EI and EII, were initially purified from extracts of germinating barley, and the genes were identified by determining the amino acid sequence of the isolated proteins (Woodward and Fincher, 1982; Woodward et al., 1982). These two isoenzymes showed 92% sequence identity at both amino acid and nucleotide levels. The gene encoding EII is expressed in the aleurone layer of germinating seeds, while the EI gene is transcribed in both germinating seeds, roots, and leaves (Slakeski et al., 1990; Slakeski and Fincher, 1992). Lichenase hydrolyzes MLG in the cell walls during seed germination, and likely both EI and EII are involved in this process to support seedling growth by producing glucose from MLG stored in the endosperm. Isoenzyme EI is also expressed in young roots and leaves. MLG content decreases in elongating coleoptiles and leaves during cell elongation. It was believed that lichenase is responsible for the removal of this MLG; however, the EI transcript was not detected in barley coleoptiles, leading to doubt as to whether EI mediates wall hydrolysis in these tissues (Slakeski and Fincher, 1992). So far, the function of EI in elongating vegetative tissues is not fully understood.

MLG amounts vary in different tissues and developmental stages with the largest amounts found in endosperm and expanding vegetative tissues. In *B. distachyon*, the endosperm contains more than 40% (w/w) MLG but less than 7% (w/w) starch which indicates a more prominent role for MLG in providing glucose for the developing embryo of germinating seeds than starch (Guillon et al., 2011). Genes encoding MLG hydrolase must exist to meet the nutrient requirement of seed germination, but so far, the lichenase responsible for the hydrolysis of MLG in the endosperm during germination have not been identified in *B. distachyon*. MLG is also highly abundant in elongating coleoptiles followed by extensive removal when cell elongation ceases. MLG accumulation is associated with cell elongation in vegetative tissues in grasses (Kim et al., 2000; Carpita et al., 2001). Walls of elongating leaves contain as high as 10% (w/w) MLG, while senesced leaves have less than 0.5% (w/w) MLG (Fan et al., 2018). Likely the majority of the MLG deposited in young leaves is removed by lichenase after cell elongation. The significance of this removal to changes in wall structure and function is not clear.

Besides providing glucose during seed germination, MLG could serve as an energy source in the dark to support plant growth as does starch. Roulin et al. reported that transcription of the EI gene was induced in barley seedling leaves after dark treatment, while no mRNA of isoenzyme EII was detected. MLG content in leaves of dark-incubated plants decreased, which is consistent with the induction of EI by darkness (Roulin et al., 2002). Recently Kraemer et al. identified a lichenase in maize named MLGH1, and this enzyme is involved in MLG degradation during the night. The mutant *mlgh1* contained more MLG and showed no significant growth defects (Kraemer et al., 2021). But so far, it is not clear how plants coordinate the hydrolysis of MLG and starch to maintain plant growth.

In many land plants, transitory starch is a primary storage product of photosynthesis and the main energy source during the night (Zeeman et al., 2004). Currently, both synthesis and degradation pathways of starch are well understood, while less is known about the regulation of these processes in response to changes in other carbohydrates such as MLG. The amount of transitory starch differs not only between day and night, but among different tissues and tissue ages. In rice, starch content decreases during the tillering stage and then increases when all the tillers are nutritionally independent (Sato, 1984). Starch metabolism is likely different during developmental stages in response to the rate of plant growth.

Here we report that a gene encoding a lichenase in *B. distachyon* is expressed in the endosperm of germinating seeds and vegetative tissues. Disruption of this gene alters the metabolism of both MLG and starch. Immunolabeling with a MLG monoclonal antibody demonstrated that in tissues that express *BdLCH1*, the MLG in plants with a disrupted *BdLCH1* gene was not removed as was the case in wild-type plants. The results provide an attractive strategy for biofuel production by modifying lichenase in bioenergy crops and also clues for uncovering the function of MLG during cell expansion.

## RESULTS

### BdLCH1, a Lichenase in *B. distachyon*

To examine the behavior of lichenase genes in *B. distachyon* and their relationship to other grass homologous genes, the coding sequences of lichenase-related genes from six grass species were obtained from Phytozome 12 (Goodstein et al., 2012; http://phytozome.jgi.doe.gov/). Phylogenetic analysis of these 24 nucleotide sequences resulted in two clades. Five out of six *B. distachyon* lichenase related genes fall into the same clade as genes encoding two barley lichenases EI and EII. Bradi2g27140 hitherto named *BdLCH1* was the most similar *B. distachyon* gene to HORVU1Hr1G057680 (EI) and HORVU7Hr1G120450 (EII) suggesting that BdLCH1 is a lichenase responsible for removing MLG (Figure 1). Amino acid sequence alignment of BdLCH1, EI, and EII showed that BdLCH1 has a higher sequence identity with EI that is expressed in both vegetative tissues and seeds (Supplemental Figure 1) (Slakeski et al., 1990; Slakeski and Fincher, 1992).

**Figure 1.**
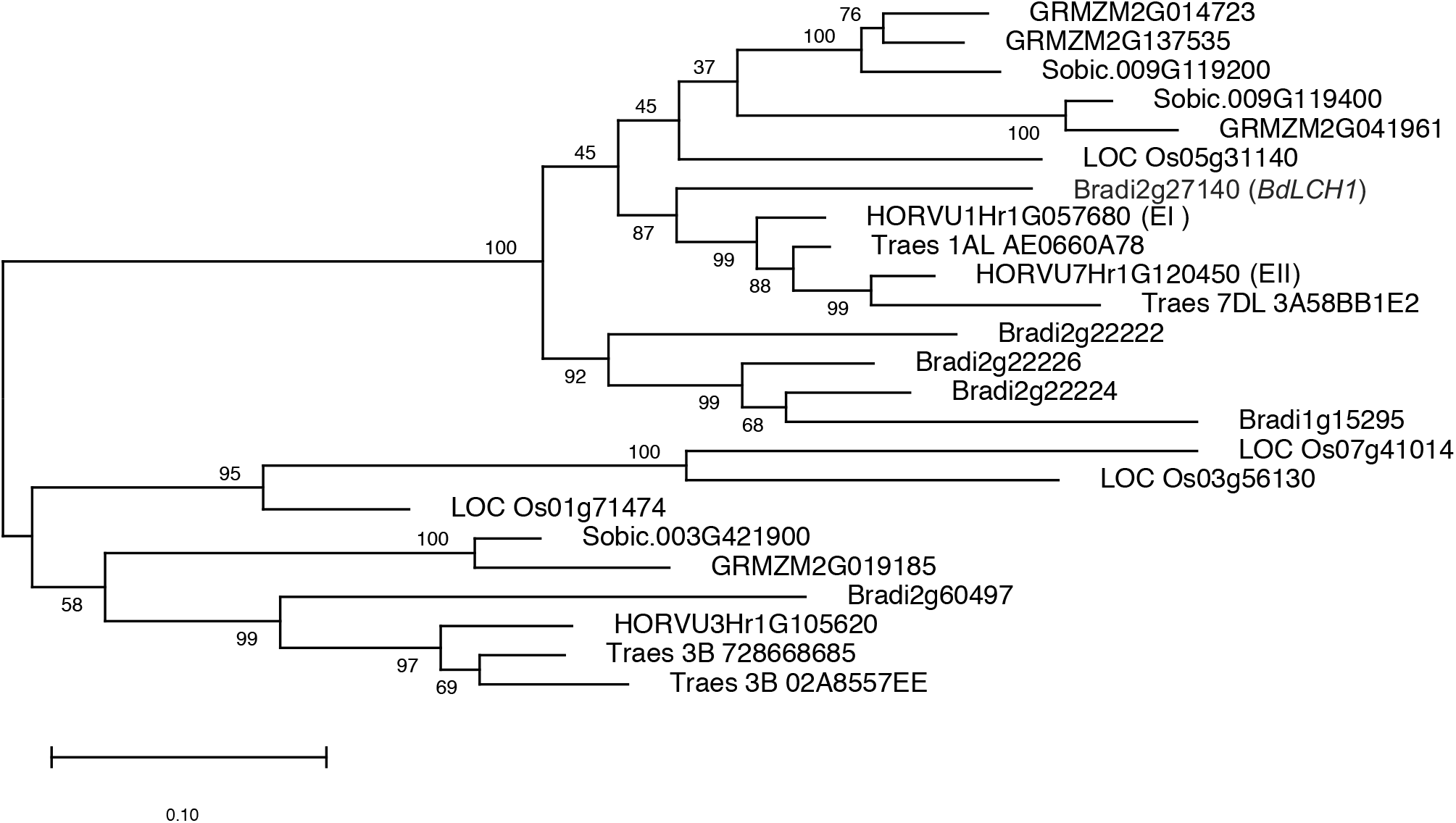
Phylogenetic Analysis of Genes Encoding Lichenase in Six Grass Species. Genes encoding putative lichenase in *Triticum aestivum, Hordeum vulgare, Brachypodium distachyon, Sorghum bicolor, Oryza sativa, and Zea mays* were analyzed using MEGA X by the Maximum Likelihood method and General Time Reversible model to generate a phylogenetic tree. The percentage of trees in which the associated taxa clustered together is shown next to the branches. The tree was drawn to scale, with branch lengths measured in the number of substitutions per site.

In order to establish that BdLCH1 has lichenase activity, we expressed BdLCH1 (amino acids 29 to 334) without the predicted N-terminal signal peptide and with a polyhistidine tag in *Escherichia coli*. Lichenase activity of the purified protein preparations was measured. The pH optima of the lichenase activity was determined as 5, which is the same as barley lichenase (Supplemental Figure 2A) (Woodward and Fincher, 1982).

The optimum temperature of BdLCH1 was found to be 30 °C (Supplemental Figure 2B). The BdLCH1 expressed in *Escherichia coli* was very active with a turnover number of 179 sec^-1^ at the maximal amount of substrate available. *K*_m_ and *V*_max_ values were not obtained due to the limitation of substrate concentration (Supplemental Figure 2C). These results establish BdLCH1 as a lichenase capable of MLG hydrolysis in vitro.

### *BdLCH1* Is Expressed in Both Grains and Vegetative Tissues

To further characterize the six putative lichenase genes of *B. distachyon*, we examined their expression in a number of tissues and developmental stages. *BdLCH1*, Bradi2g22222, Bradi2g22224, and Bradi2g22226 are expressed at very high levels in the endosperm of germinating seeds. *BdLCH1* is also transcribed at a relatively high level in elongating internodes and at low levels in fully elongated internodes and leaves (Figure 2A). To obtain a developmental profile with high resolution, we dissected elongating internodes (EI) in eight successive 1 mm pieces (EI-1 to 8) followed by two 2 mm pieces (EI-9 and 10) for RNA profiling. EI-1 is located at the bottom of the elongating internode and EI-10 is at the top of the elongating internode. To gain information to determine the cell developmental stage of each section, we examined expression levels of *BdCESA1* and *BdCSLF6*, which are associated with primary cell wall formation and MLG synthesis, respectively. The expression of both genes was highest at stage EI-3 and then decreased with following stages indicating that cells in EI-3 underwent active cell elongation and the elongation slowed down in EI-4 and 5 (Supplemental Figure 3A and 3B). *BdLCH1* had the highest expression level in EI-5 indicating that it was expressed after the cell elongation phase. To further examine the expression pattern of *BdLCH1*, 24 h, 48 h, 72 h, and 120 h coleoptiles were examined. *BdCSLF6* and *BdCESA1* were most highly expressed in 24 h coleoptiles followed by a rapid decrease indicating cells in 24 h coleoptiles were at a stage of rapid elongation. On the contrary, transcripts of *BdLCH1* were first detected in 48 h coleoptiles when elongation had ceased, with expression increasing rapidly in following stages. Transcription of *BdLCH1* was not detected in 24 h coleoptiles. In 48 h coleoptiles, the expression level of *BdCSLF6* began to decrease while the transcription of *BdLCH1* increased rapidly in 72 h and 120 h coleoptiles (Figure 2B). These results are consistent with a previous report in maize coleoptiles that MLG is hydrolyzed when cell elongation and synthesis of MLG ceases (Kim et al., 2000) and indicate that BdLCH1 is responsible for the turnover of MLG after cell elongation cessation. No expression of Bradi2g22222, Bradi2g22224 or Bradi2g22226 was seen in the examined vegetative tissues which suggests a specific function of these proteins during seed germination (Figure 2A). Two of the six *B. distachyon* lichenase related genes, Bradi1g15295 and Bradi2g60497, were expressed at a very low level in some of the examined vegetative tissues but were not expressed in endosperm of germinating seeds (Supplemental Figure 3C). Hence, *BdLCH1* is the dominant lichenase expressed in vegetative tissues, including coleoptiles, internodes, and leaves, while also being one out of four highly expressed homologs in germinating endosperms.

**Figure 2.**
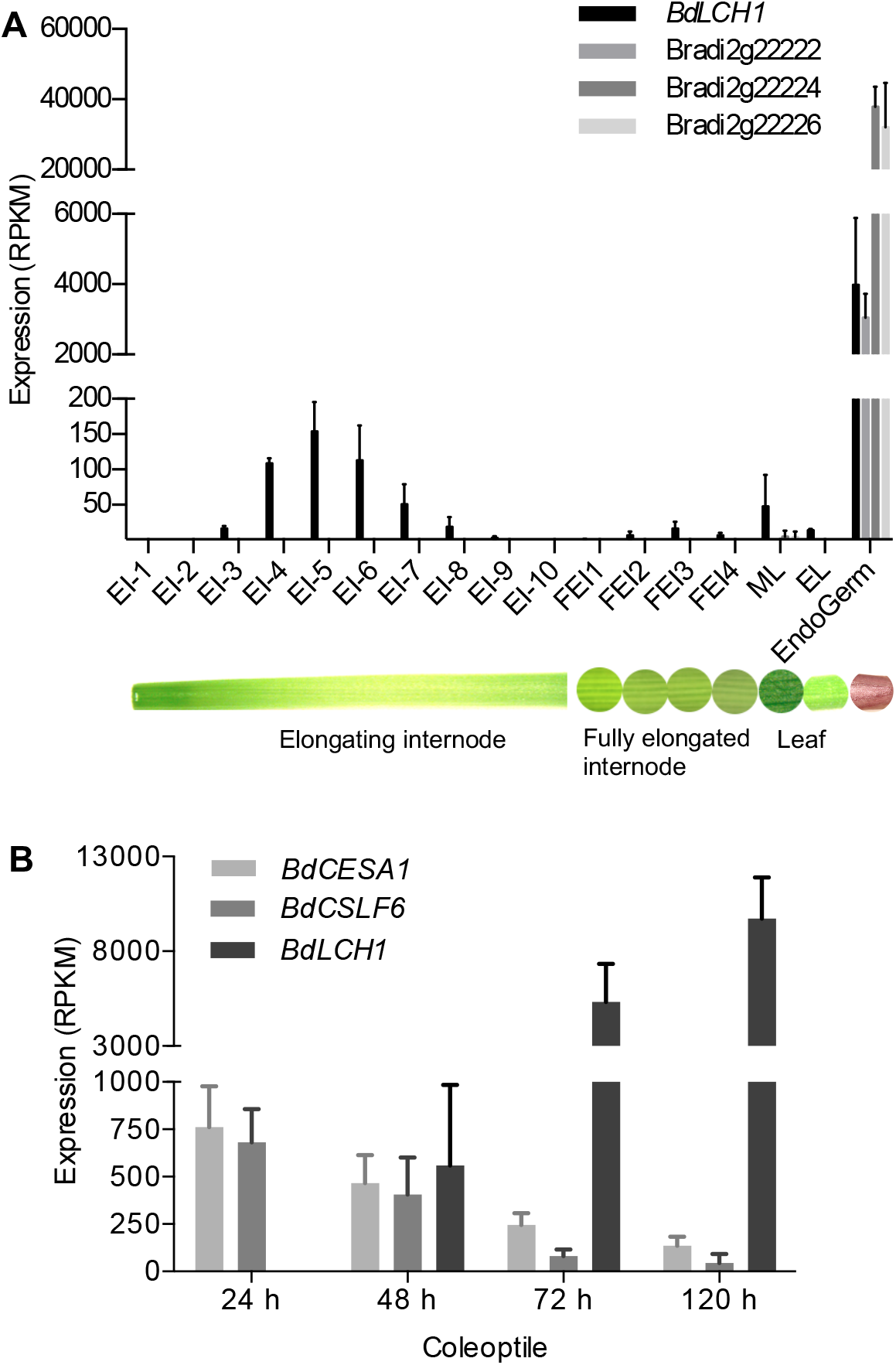
Expression Profiles of Four Lichenase Genes, *BdCESA1*, and *BdCSLF6* in *Brachypodium distachyon*. **(A)** Tissue samples from *B. distachyon* plants grown under stem elongation conditions (16-h-light/8-h-dark cycle) were used to prepare RNA for expression analysis. EI-1 to EI-10 indicate 10 successive segments from elongating internodes of nine-week-old plants. Samples EI-1 to EI-8 are 1 mm long and samples EI-9 and EI-10 are 2 mm long. EI-1 is located at the base of elongating internodes and EI-10 is at the top of elongating internodes. FEI1 to FEI4 indicate samples from fully elongated internodes from 12-week-old plants. FEI1 and FEI4 are the youngest and oldest internodes, respectively. ML, mature leaf blades from 12-week-old plants. EL, the base of elongating leaf that were fully enclosed by the sheath of the next older leaf. EndoGerm, the endosperm of germinating seeds. **(B)** Expression profiles of *BdCESA1, BdLCH1*, and *BdCSLF6* in coleoptiles. Coleoptiles for RNA-seq were harvested from dark-grown seedlings when they were 24 h, 48 h, 72 h, or 120 h old. RPKM, Reads Per Kilobase of transcript per Million mapped reads. Expression is shown as means ± sd from three biological replicates.

To determine the correlation between the expression pattern of *BdLCH1* and MLG degradation at a cellular level, we examined the expression of *BdLCH1* in mature vegetative tissues using in situ hybridization and MLG deposition by immunolabeling with an anti-MLG monoclonal antibody. *BdLCH1* expression was detected in chlorenchyma cells of fully elongated internodes (Figure 3A) and fully expanded leaves (Figure 3B). *B. distachyon* young internodes and leaves accumulate large amounts of MLG in those cells (Supplemental Figure 4), while no MLG was detected in these cell types in fully expanded internodes and leaves (Figure 5, Supplemental Figure 6B) when cross sections were labeled with MLG antibody. These results indicate that MLG degradation correlates with the expression of *BdLCH1* in internodes and leaves. MLG in the cells expressing *BdLCH1* was degraded after cell elongation, which indicates that BdLCH1 is responsible for removing MLG in chlorenchyma cells of internodes and leaves during plant development.

**Figure 3.**
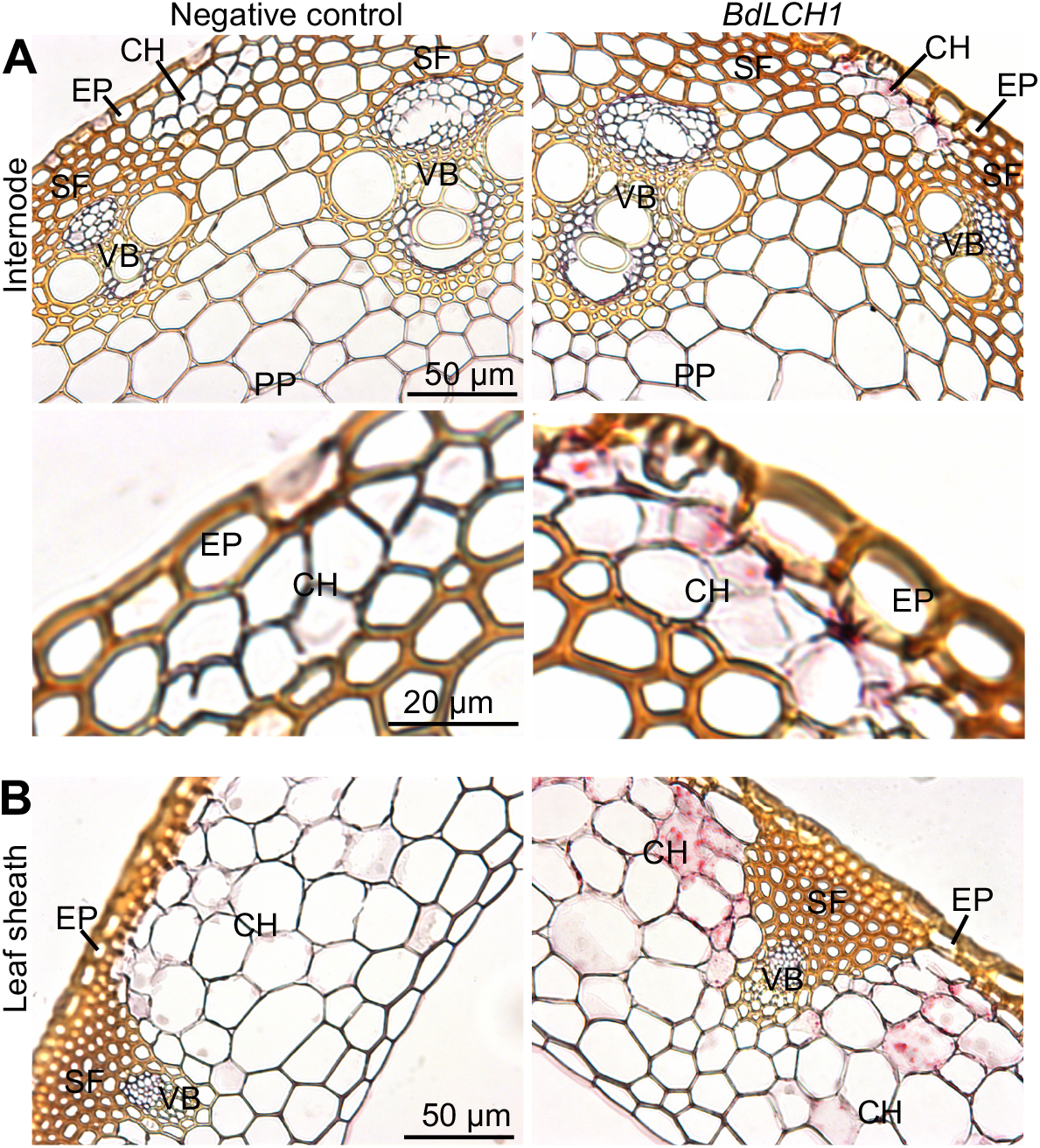
Cellular Localization of *BdLCH1* in Mature Vegetative Tissues. **(A)** and **(B)** In situ hybridization of *BdLCH1* in fully elongated internodes **(A)** and mature leaf sheaths **(B)**. Probes that target the bacterial *DapB* gene were used as negative control. Each red punctuate dot in the RNAscope assay represents a single mRNA transcript. Bottom panel in **(A)** shows an enlarge image of a region with chlorenchyma cells of fully elongated internodes (top panel). CH, chlorenchyma cell; EP, epidermis cell; PP, pith parenchyma cell; SF, sclerenchyma fiber; VB, vascular bundle.

### *BdLCH1* Mutants Accumulate More MLG in Both Young and Senesced Vegetative Tissues

To discover the role of *BdLCH1* on MLG accumulation, we generated *lch1* mutants using the CRISPR-Cas9 gene editing system (Xie et al., 2015). The construct used to induce gene lesions included two guide RNAs (gRNAs) separated by 129 nucleotides. Six independent homozygous lines were obtained and two of them, namely *lch1-c1* and *lch1-c2*, were examined further. Sequencing results showed that *lch1-c1* and *lch1-c2* have eight and 11 nucleotides deletions located 472 nucleotides downstream of the start codon, respectively (Supplemental Figure 5). This resulted in frameshifts in both mutants leading to premature termination codons in both cases, which significantly truncates the wild-type BdLCH1 protein (165 amino acids compared to 334 amino acids of wild type). *lch1-c1* was 18% shorter in height and 15% smaller in stem diameter than wild-type plants, while such differences were not observed for *lch1-c2* (Figure 4A-C). Both mutants showed no significant difference in vegetative plant dry weight (Figure 4D). These results indicate that disruption of *BdLCH1* may not necessarily affect plant growth.

**Figure 4.**
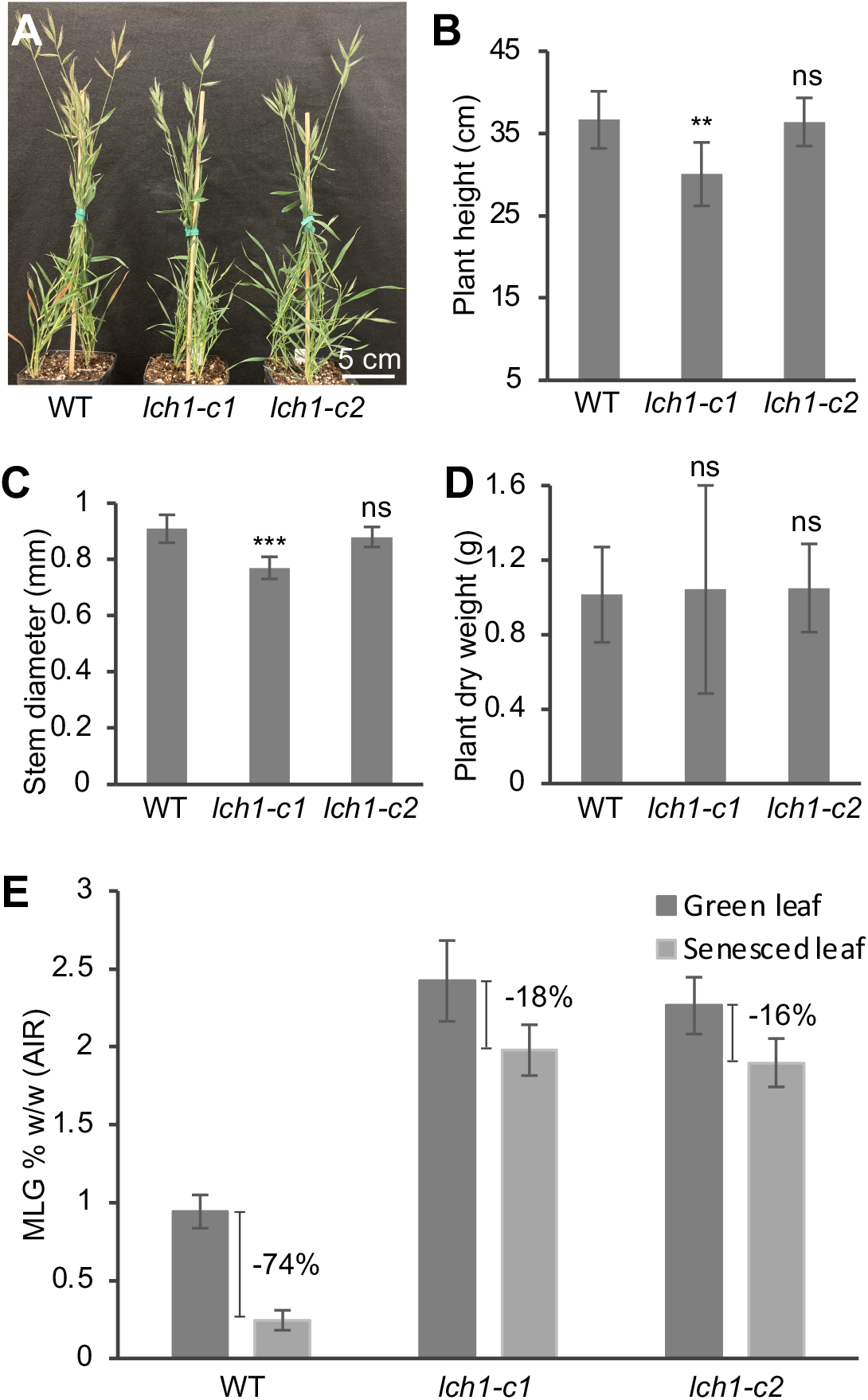
Growth Measurements and MLG Content of *BdLCH1* CRISPR Lines. **(A)** Phenotype of 10-week-old wild type (WT) and two independent *BdLCH1* CRISPR lines grown under long day condition (20-h-light/4-h-dark). **(B)** to **(D)**, Measurements of plant height **(B)**, stem diameter **(C)**, and vegetative plant dry weight of senesced plants **(D)**. Measurements are shown as means ± sd from six biological replicates. Asterisks indicate statistically significant differences at P < 0.01 (**) and P < 0.001 (***) using Student’s *t* test. ns indicates no statistically significant difference. **(E)** MLG amounts in green leaves of eight-week-old plants and senesced leaves are shown as means ± sd from five biological replicates. Alcohol insoluble residues (AIR) extracted from leaves were subjected to MLG assays.

Analysis of total MLG content of alcohol-insoluble residues (AIR) prepared from mature green leaves showed that these mutants contained approximately twice the amount of MLG compared to wild type, i.e. 2.2% (w/w) MLG in mutants compared to 0.9% (w/w) MLG in wild-type plants (Figure 4E). In senesced leaves, wild-type plants contained 0.2% (w/w) MLG while both mutants contained 1.9% (w/w) MLG. This is consistent with *BdLCH1* being expressed in chlorenchyma cells of mature leaves (Figure 3B), which indicates MLG turnover occurs in green mature leaves. Hence, MLG content was reduced by 74% upon senescence in the wild type and less than 20% in the mutants. The combination of increased MLG accumulation in mature green leaves and reduced turnover during senescence brings the final MLG content in the senesced leaves to an eight-fold higher in the mutants compared to wild-type plants (Figure 4E). Given this result we also investigated MLG content of senescent internodes where we found a similar pattern of four-fold higher MLG content in the mutants, i.e. 0.8% (w/w) MLG in the mutants compared to 0.2% (w/w) in wild-type plants (Supplemental Figure 6A). We conclude that MLG accumulation is increased and degradation is reduced in *lch1* mutants, while no apparent growth defect was observed.

### Abundance of MLG in Chlorenchyma Cells of *lch1* Mature Tissues

To determine MLG accumulation in *lch1* mutants at the cellular level, cross sections of mature leaf blades and leaf sheaths were labeled with MLG antibody. In leaf sheaths, a strong signal was present in vascular bundles, sclerenchyma cells, and epidermal cells in both wild-type and mutant plants. A strong signal was also detected in chlorenchyma cells of *lch1* mutants which indicates MLG accumulation in these cells, while no obvious binding of the antibody was observed in mature chlorenchyma cells of wild-type plants (Figure 5A). This result is consistent with the in situ hybridization data showing that *BdLCH1* is expressed in chlorenchyma cells in wild type (Figure 3B) and implicates BdLCH1 in MLG hydrolysis in this cell type, i.e. without a functional BdLCH1, MLG was stabilized in chlorenchyma cells.

**Figure 5.**
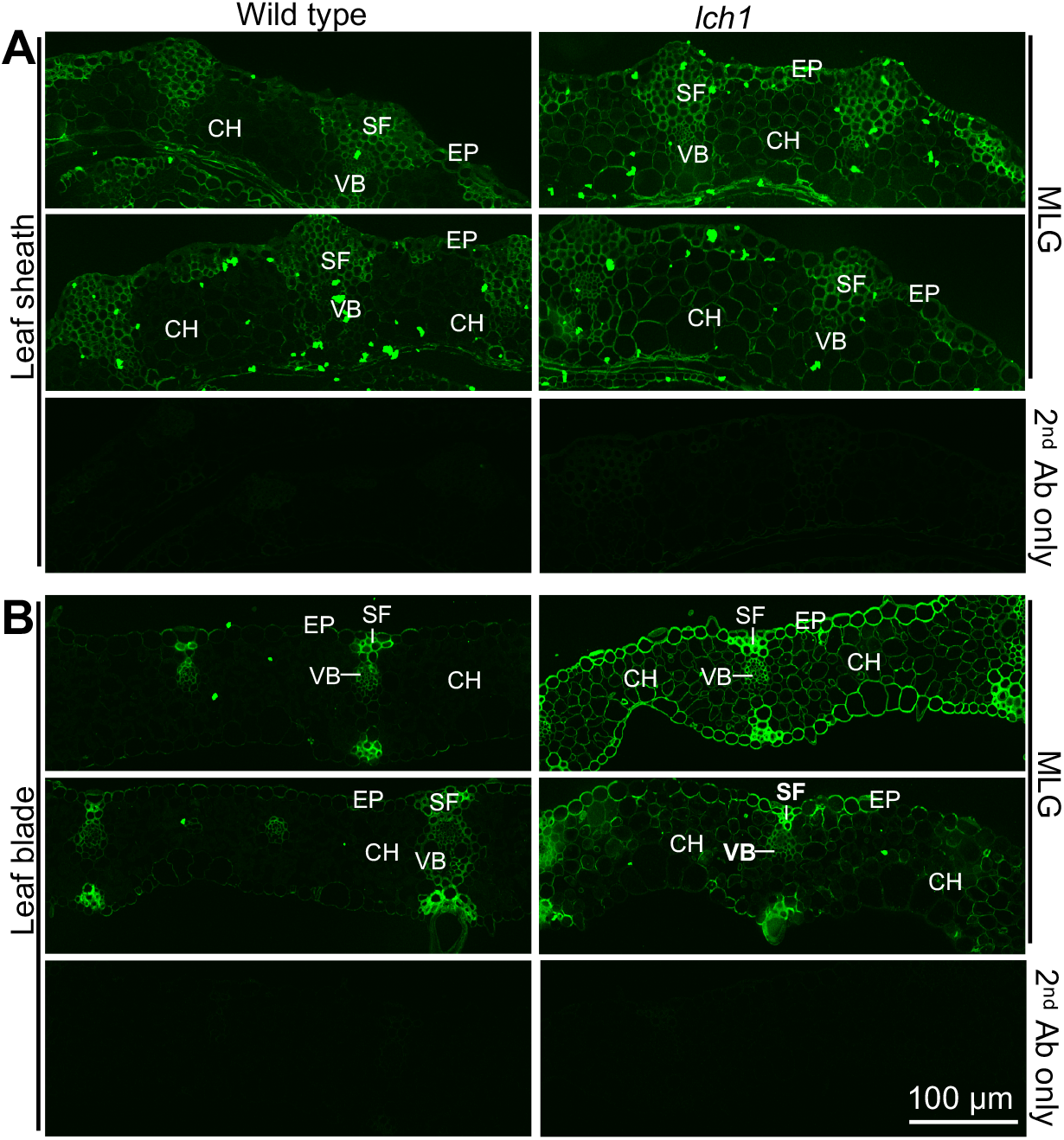
Immunolabeling of Leaf Sheath and Leaf Blade Sections with MLG Monoclonal Antibody. **(A)** and **(B)** Sections from leaf sheaths **(A)** and leaf blades **(B)** of two biological replicates of wild type (left panel) and two independent *lch1* lines (right panel) were immunolabeled with MLG monoclonal antibody as shown in the top two panels of **(A)** and **(B)**. Sections immunolabeled with only secondary antibody were used as negative control. EP, epidermis cell; SF, sclerenchyma fiber; VB, vascular bundle.

Similar observations were made in leaf blades. MLG was also presented in chlorenchyma cells of *lch1* mutants but not in wild-type plants. There was strong labeling in vascular bundles and sclerenchyma cells in both wild type and mutants. A strong signal was also observed in epidermal cells of *lch1* plants while only weak labeling was observed in wild-type plants (Figure 5B). No signal was detected in negative controls (secondary antibody only) for both leaf sheaths and leaf blades (Figure 5).

We also examined MLG deposition in mature internodes by immunolabeling. Binding of the MLG antibody was detected in all cells examined with the exception of chlorenchyma cells in wild-type plant samples while weak labeling was detected in this cell type of *lch1* plants (Supplemental Figure 6B). Notably, the in vitro hybridization data showed *BdLCH1* expression in chlorenchyma cells of fully elongated internodes (Figure 3A). Hence, these results implicate BdLCH1 in the removal of MLG in chlorenchyma cells of mature internodes. To further confirm MLG deposition at cellular level, a MLG antibody (Meikle *et al*., 1994) was used for immunogold labelling of fully elongated internodes. Gold particles were found in walls of all cells but the labeling density was different depending on cell types. Consistent with the immunofluorescence labeling results, walls of epidermis, sclerenchyma, and pith parenchyma cells were heavily labeled while few gold particles were present in walls of chlorenchyma cells (Supplemental Figure 7). Taken together, the immunolabeling results from mature leaf and internode samples confirmed that *lch1* mutants accumulate more MLG than wild-type plants and implicate the BdLCH1 enzyme in removing MLG in chlorenchyma cells in mature vegetative tissues.

### Transcription of *BdLCH1* Is Induced in Leaves of Dark-incubated *B. distachyon*

Barley isoenzyme EI was induced in dark-incubated barley seedlings and MLG content decreased significantly (Roulin et al., 2002). Similar result was found in maize, expression of *MLGH1*, a gene encoding lichenase, was increased during the night (Kraemer et al., 2021). To determine whether genes encoding lichenases in *B. distachyon* are induced by darkness and to further examine the response of *lch1* mutants to darkness, three-week-old wild-type and *lch1* plants were incubated under a 16-h-light/8-h-dark cycle (0 h), or in complete darkness for 24 h, 48 h, and 72 h (Supplemental Figure 8). Reverse transcription quantitative PCR (RT-qPCR) analysis showed that *BdLCH1* was induced in leaves of dark-incubated wild-type and *lch1* mutant leaves. The expression level of *BdLCH1* was significantly lower in mutant plants than that in wild-type plants, which suggests nonsense-mediated mRNA decay (NMD) of the mutant mRNA was elicited by the premature termination codons (Figure 6A) (Peltz et al., 1993; Popp and Maquat, 2016). Induction of *lch1* mRNA in both mutants in darkness suggests the cis-regulatory element is intact in both *lch1-c1* and *c2*. These results show that *BdLCH1* is induced in leaves of dark-incubated *B. distachyon* plants.

**Figure 6.**
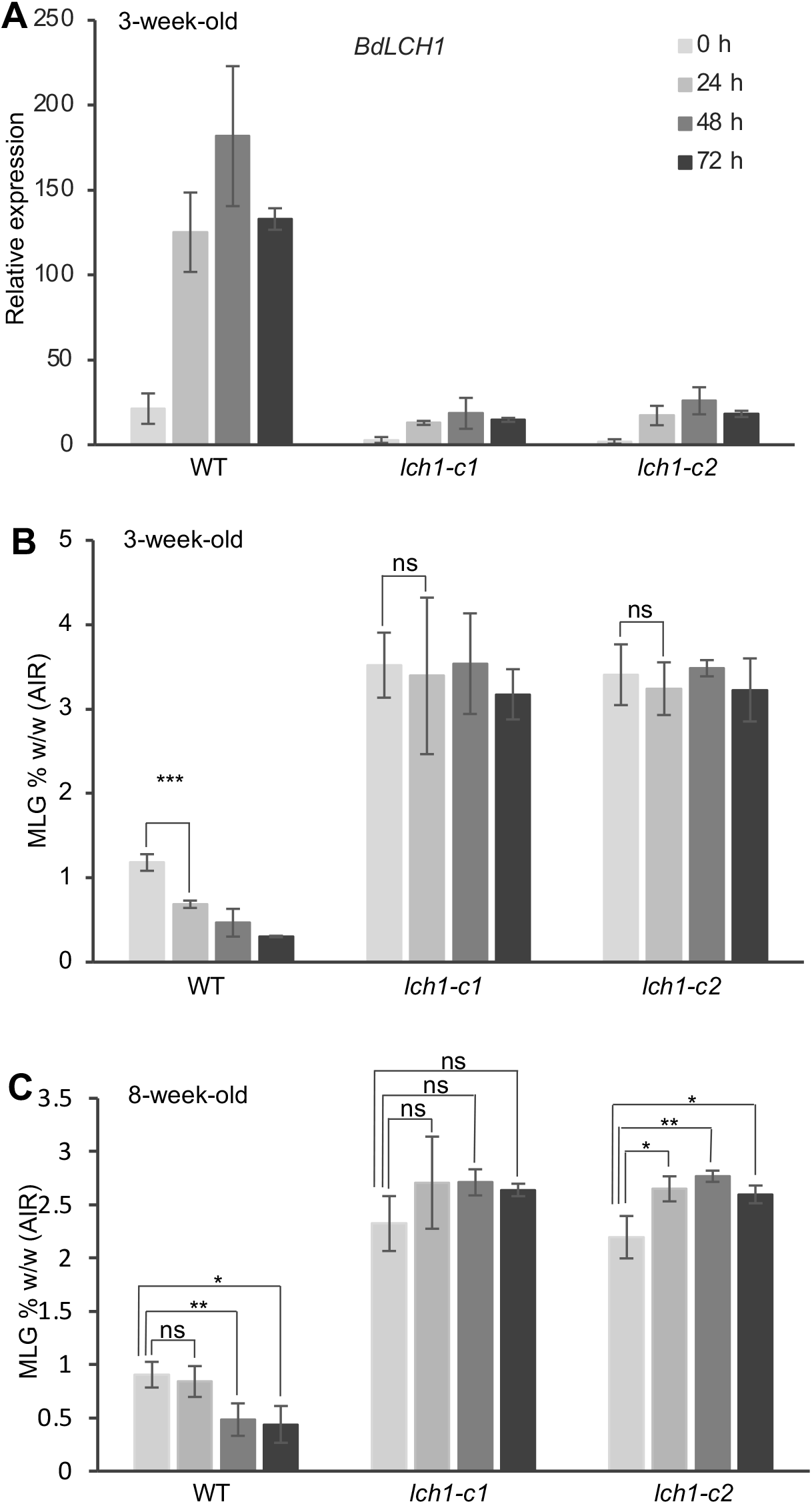
*BdLCH1* transcription Levels and MLG Content in Mature Leaves of Dark-incubated Plants. **(A)** RT-qPCR analysis of *BdLCH1* expression levels in dark-incubated wild type (WT) and *lch1* lines. Three-week-old plants were incubated under regular light/dark cycle (0 h), or in complete darkness for 24 h, 48 h, and 72 h. Relative expression levels are presented as means ± sd of three biological replicates. Dark-incubation and sampling of the leaves were started in the middle of the light period. **(B)** and **(C)** MLG amounts in three-week-old **(B)** or eight-week-old plants **(C)**. Plants were incubated under regular light/dark cycle (0 h), or in complete darkness for 24 h, 48 h, and 72 h. Dark-incubation and sampling of the leaves were started in the middle of the light period. Alcohol insoluble residues (AIR) extracted from leaves were assayed for MLG content. MLG amounts are shown as means ± sd from three **(B)** or two **(C)** biological replicates. Asterisks indicate statistically significant differences at P < 0.05 (*), P < 0.01 (**), and P < 0.001 (***) using Student’s *t* test. ns indicates no statistically significant difference.

### MLG Accumulation in *lch1* Mutants Is Resistant to Dark Induced Degradation

To investigate MLG accumulation in dark-incubated plants, we determined total MLG content in AIR prepared from mature leaves of three-week-old plants placed in the dark for 0, 24, 48, and 72 h. MLG amounts in three-week-old wild-type plants decreased significantly after dark-incubation for 24 h and dropped further to 26% of the starting levels after 72 h dark-incubation, while leaf MLG content of dark-incubated mutant plants was similar when grown under 0, 24, 48, or 72 h of darkness (Figure 6B). This result implicates BdLCH1 in the removal of MLG in leaves of dark grown plants.

To understand the regulation of MLG biosynthesis in leaves of dark-incubated plants, expression levels of *BdCSLF6*, which encodes the primary MLG synthase, were determined using RT-qPCR. The result shows that *lch1* plants had similar expression levels of *BdCSLF6* as that of wild-type plants. Expression level of *BdCSLF6* decreased in both wild-type and mutant plants after dark incubation (Supplemental Figure 9). This result shows that MLG biosynthesis, as indicated by *BdCSLF6* expression, was blocked in darkness in both wild type and *lch1* plants.

To examine MLG content and response to darkness of older plants, eight-week-old wild type and *lch1* mutants were incubated in the dark for 0, 24, 48, and 72 h. Measurements of the MLG content of mature leaves of eight-week-old plants showed that MLG degradation in wild type was not as fast as that in three-week-old plants as MLG content was not significantly decreased after 24 h of dark incubation, while 51% of MLG was hydrolyzed after 72 h of dark incubation (Figure 6B and 6C). This suggests that the induction of lichenase in older plants is lower in response to darkness than in younger plants. Similar to three-week-old plants, MLG accumulation in dark-incubated *lch1* mutants was not reduced compared to mutants grown under 16-h-light/8-h-dark cycle (0 h), in fact, *lch1* mutants accumulated more MLG during the dark-incubation (Figure 6C). Dark-incubation of three-week-old and eight-week-old plants indicates that *lch1* mutant plants do not degrade MLG in response to darkness and accumulate more MLG than wild type in both young and old plants.

### Disruption of BdLCH1 Alters Starch Degradation in Darkness

Starch is the main storage carbohydrate in most grasses and is often degraded at night to sustain metabolism and growth. To examine starch content, AIR prepared from mature leaves of three-week-old and eight-week-old plants that had been placed in the dark for 0, 24, 48, and 72 h were analyzed for starch content. Starch amounts in leaves of three-week-old plants of both wild type and *lch1* decreased markedly after dark-incubation to background levels (more than 95% reduction) after 24 h and remained at that level for the two later time points (Figure 7A). In eight-week-old plants, mature leaves of *lch1* and wild-type plants accumulate a similar amount of starch under standard growth conditions, but the amount of starch observed under dark incubation was lower in mutant plants than in wild-type plants. After 24 h of dark incubation, 37% of the starch was degraded in wild-type plants, as opposed to 85% and 89% in *lch1-c1* and *lch1-c2*, respectively. Starch content continued to decrease after 48 and 72 h of continuous darkness (Figure 7B). These results indicate that the *lch1* mutants showed a different response to darkness with respect to starch metabolism as compared to the eight-week-old wild-type plants. To confirm these results, starch in leaf blades and leaf sheaths was visualized using Lugol’s iodine staining reagent. No difference was observed between wild type and mutants when grown under 16-h-light/8-h-dark cycle (Figure 7C and 7D). When plants were grown under 24 h of continuous darkness, three-week-old *lch1* mutant showed similar staining as wild type in both leaf blades and leaf sheaths (Figure 7C). However, staining of leaf blades and leaf sheaths of eight-week-old *lch1* mutant plants revealed a lower amount of starch in comparison to wild-type plants (Figure 7D). To examine starch deposition at cellular level, cross sections of internodes and leaves were stained. Staining signal was present in chlorenchyma cells in internode and leaf tissues (Supplemental Figure 10). We conclude based on both total starch quantification and in situ staining, that the *lch1* mutants degrade starch under dark treatment at a faster rate in older plants.

**Figure 7.**
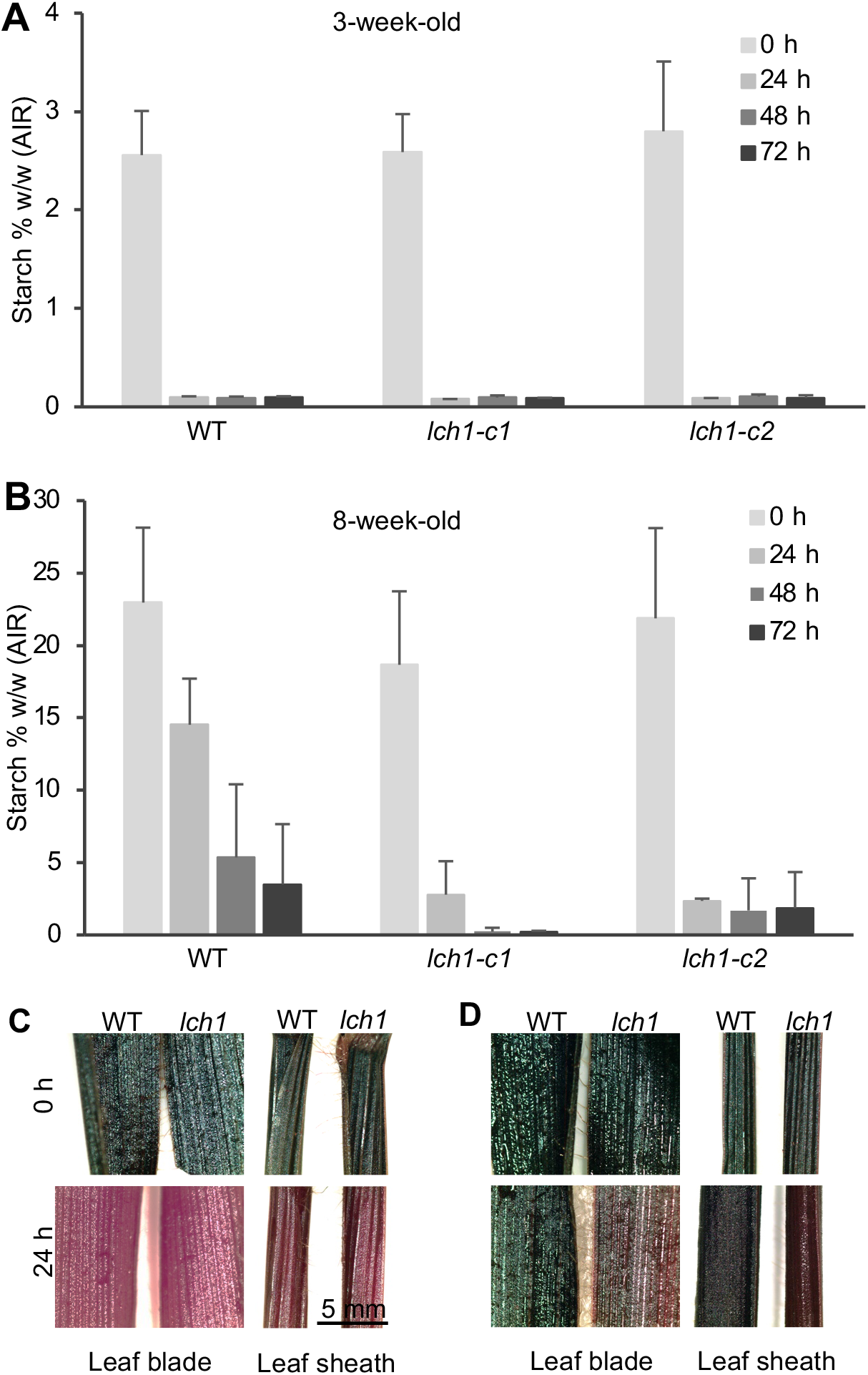
Starch Content and Iodine Staining of Mature Leaves of Dark-incubated Plants. **(A)** and **(B)** Starch amounts in mature leaves of dark-incubated three-week-old **(A)** or eight-week-old plants **(B)**. Plants were incubated under regular light/dark cycle (0 h), or in complete darkness for 24 h, 48 h, and 72 h. Dark-incubation and sampling of the leaves were started in the middle of the light period. Alcohol insoluble residues (AIR) extracted from leaves were subjected to starch assays. Starch amounts are shown as means ± sd from three **(A)** or two **(B)** biological replicates. **(C)** and **(D)** Iodine staining of leaf blades (left panel) and leaf sheaths (right panel) of three-week-old **(C)** or eight-week-old plants **(D)**. Tillers were cut at the bottom and incubated in culture tubes with distilled water under regular light/dark cycle (0 h), or in complete darkness for 24 h.

## DISCUSSION

### MLG Removal During Cell Expansion

In vegetative tissues, MLG is a developmental stage-specific polysaccharide that is synthesized and degraded when cell expansion begins and ends, respectively. Elongating vegetative tissues showed the greatest content of MLG when cells were actively expanding (Kim et al., 2000; Carpita et al., 2001; Gibeaut et al., 2005). Our transcriptome analysis of coleoptiles showed that expression of *BdLCH1* was not detected in 24 h coleoptiles which underwent active cell elongation. The expression was first detected in 48 h coleoptiles and increased in 72 h and 120 h coleoptiles (Figure 2B). Cell elongation began to slow down in coleoptiles of 48 h old and 72 h coleoptiles which were fully expanded. Together with previous studies, these results indicate that BdLCH1 is responsible for removing MLG in fully expanded cells. As was seen in coleoptiles, the expression level of *BdLCH1* in different sections of elongating internode is correlated with cell elongation. *BdLCH1* showed the highest transcription level in EI-5 that cell expansion ceased (Figure 2A). MLG removal was also observed at cellular level by immunolabeling with MLG antibody. MLG was ubiquitous in elongating leaves and internodes (Supplemental Figure 4), while removed in chlorenchyma cells after cell elongation (Figure 5; Supplemental Figure 6B and 7). The majority of MLG in chlorenchyma cells was degraded after cell elongation, while there are still detectable level of MLG in other cell types. Epidermis cells, sclerenchyma cells, and cells in the vascular bundle still contained high levels of MLG (Supplemental Figure 7). These cells have thick lignified walls with MLG. Other mature cells, such as pith parenchyma cells, which have primary walls, also retained MLG. However, the MLG in chlorenchyma cells in both leaves and internodes was hydrolyzed after cell elongation was completed (Figure 5 and Supplemental Figure 7). The cell type removal of MLG correlates with the expression pattern of *BdLCH1*. The significance of removing MLG from vegetative tissues when cell elongation begins to slow is not clear. Our hypothesis is that MLG is detrimental to the functioning of the mature cell wall in certain cell types such as chlorenchyma. The cell types that express the *BdLCH1* gene are those that are highly metabolically active which suggests that removal of MLG may affect the permeability of the wall to allow higher flux of metabolic substrates such as CO_2_ (Ellsworth et al., 2018; Liu et al., 2019). However, we did not observe significant growth defects in *lch1* mutants and a similar result was reported in maize *mlgh1* mutants (Kraemer et al., 2021). The plants in our study were grown under controlled growth conditions and as such we do not know if there would be an effect of the lichenase deletion on plant development under stress conditions.

### Starch Metabolism of *lch1* Mutants

Transitory starch is the major storage carbohydrate synthesized during the day in higher plants and is broken down to sustain plant growth at night. Darkness during the day can also trigger starch degradation (Zeeman et al., 2002). In this study, both wild type and *lch1* mutants showed a response to darkness on starch degradation and starch level decreased after 24 h dark-incubation, with eight-week-old *lch1* plants showing a faster rate of starch breakdown than wild-type plants (Figure 7B and 7D). MLG in barley leaves was also suggested to be a transitory storage carbohydrate made during the day and used as an alternative energy source to support growth at night (Roulin et al., 2002), while we find this is also the case for *B. distachyon*. A possible explanation for the faster starch degradation in *lch1* mutants is that impaired MLG hydrolysis requires additional starch degradation to meet the metabolic energy needs of plant growth in the dark. Countering this argument is the fact that the MLG levels are much lower than the starch levels. In eight-week-old plants, there was more than 20% (w/w) of starch and only 0.9% and 2.3 % (w/w) of MLG in wild-type and *lch1* plants, respectively (Figure 6C and 7B). The regulation of these two stores of fixed carbon may be linked but the connection is not likely due to the loss of available glucose from the MLG pool. It is more likely that MLG or BdLCH1 affects the signal transduction network involved in the regulation of starch metabolism. We only observed a difference in starch metabolism between wild type and *lch1* in eight-week-old plants but not in three-week-old plants.

Starch stores are approximately 10 times less in the three-week-old plants and depleted within the first 24 h of dark treatment (Figure 7A and 7C). The difference in starch breakdown between three-week-old wild-type and *lch1* mutant plants may occur in the first few hours after dark incubation and if so would not have been detected in our study. Starch content is variable in relation to plant growth (Sato, 1984). The three-week-old plants we examined were at the end of the tillering stage and the growth of younger tillers are nutritionally dependent on their mother tillers, so the amount of starch is low at this stage. The eight-week-old plants we used were at the later stem elongation stage. All tillers were able to deposit starch, so the content is higher than three-week-old plants.

Iodine staining of leaf and internode cross sections showed that chlorenchyma cells contain high levels of starch (Supplemental Figure 10), similar to previous study in *B. distachyon* (Jensen and Wilkerson, 2017). These cells have high levels of *BdLCH1* expression and MLG removal in mature cells. Chlorenchyma cells are photosynthetic and highly metabolically active. The effect of BdLCH1 on starch metabolism may indicate a role of the changes in mature wall function due to MLG removal in the regulation of starch metabolism.

### A Promising Strategy of Engineering Bioenergy Plants

Engineering bioenergy plants to accumulate a large amount of sugars for biofuels is a key strategy to substitute fossil fuels. MLG exists predominantly in cell walls of grasses. It is an easily fermentable cell wall polysaccharide and ideal compound for the production of biofuels. Increased MLG content in plants by overexpressing MLG biosynthesis genes has been achieved, but with a detrimental effect on plant growth and biomass yield (Burton et al., 2011; Vega-Sánchez et al., 2015; Kim et al., 2018; Fan et al., 2018). In this study, we identified a gene *LCH1* encoding a *B. distachyon* lichenase that likely degrades MLG in mature cell walls and studied the effect of deletion mutations in this gene on plant growth and starch metabolism. Total MLG quantification and immunolabeling using MLG antibody showed that *lch1* mutants accumulated more MLG in both leaves and stems. In senesced leaves, there was an eight-fold increase in MLG content (Figure 4E). These levels were approximately the same or significantly higher than achieved by overexpressing MLG biosynthesis genes.

Importantly, *lch1* mutants with a large amount of MLG accumulation did not show significant growth defects in our growth condition. The mutants exhibited similar plant height and vegetative biomass yields as wild type. Further, more MLG was stored in the mutants than in the wild-type due to reduced degradation. When MLG content in senesced leaves was compared with that in fully expanded leaves, 74% of the deposited MLG was degraded in wild type tissues, whereas only 18% and 16% of the initially present MLG was degraded in *lch1-c1* and *lch1-c2* mutant plants, respectively (Figure 4E). As the MLG content was measured as a percentage of the total cell wall amount, part of the decrease may be due to the increase of other cell wall polymers like lignin during plant senescence. MLG degradation during senescence poses a challenge for using crop residues for bioenergy production due to the low amount of MLG, while this is effectively elevated in *lch1* mutants. Finally, we find that the MLG content of *lch1* plants is insensitive to dark treatment. This could be useful for post-harvest of plants for bioenergy production as this material is exposed to prolonged dark periods and therefore initiates MLG degradation, which negatively affects the polymer yield for industrial applications. Modifying *LCH1* can overcome this problem and increase biofuel production efficiency. A similar result was reported in maize that the lichenase mutant *mlgh1* contained more glucose (Kraemer et al., 2021). Our study therefore suggests that engineering bioenergy crops like maize and energy sorghum by modifying *LCH1* is a promising strategy for biofuel production.

### Putative Enzymes Hydrolyzing MLG in Endosperm During Seed Germination

*B. distachyon* endosperm contains significantly more MLG than do most other cereals (Guillon et al., 2011; Burton and Fincher, 2014). The release of glucose during germination requires a lichenase. *B. distachyon* RNA-seq data analysis showed that four out of six putative lichenase genes namely *BdLCH1*, Bradi2g22222, Bradi2g22224, and Bradi2g22226, are highly expressed in the endosperm of germinating seeds. Bradi2g22222, Bradi2g22224, and Bradi2g22226 are specifically expressed in the endosperm of germinating seeds compared to the other tissues tested. Bradi2g22222 has a similar level of expression as *BdLCH1* while Bradi2g22224 and Brdi2g22226 are expressed to extraordinarily high levels in this tissue (Figure 2A). These expression data suggest that these four genes are involved in seed germination, likely hydrolyzing storage cell wall polysaccharide MLG into sugars for growth of the seedling. Additionally, phylogenetic analysis of putative genes encoding lichenases from six grass species showed that these four genes fall into the same clade (Figure 1). This result supports the hypothesis that genes in this clade are responsible for hydrolyzing MLG during seed germination in species that use MLG as storage carbohydrate. Locations of Bradi2g22222, Bradi2g22224, and Bradi2g22226 in the genome show that they are tandemly arrayed genes likely due to recent tandem duplications. It is likely that this duplication allows for increased production of lichenase during germination.

## METHODS

### Plant Materials and Growth Conditions

*Brachypodium distachyon* ecotype Bd21-3 was used in all experiments. Seeds of *B. distachyon* were sown in Jiffy 7 peat pellets (Greenhouse Megastore) and stratified at 4°C for three days. Plants were then grown in controlled growth chambers under stem elongation conditions (16-h-light/8-h-dark cycle) as previously described (Jensen and Wilkerson, 2017), or long-day conditions (20-h-light/4-h-dark cycle) for producing seeds (Fan et al., 2018). For dark-incubation experiments, treatments were started in the middle of light period (Supplemental Figure 8). Leaf tissue of freeze-dried plants was proceeded with MLG or starch assays.

### Plasmid Construction and *B. distachyon* Transformation

The coding sequence of BdLCH1 29 to 334 amino acids was synthesized and cloned into pET151 vector (Thermo Fisher Scientific) for expressing BdLCH1 with a polyhistidine tag in BL21 of *Escherichia coli*. Sequences of two guide RNAs for CRISPR-Cas9 construct were selected using the Benchling tool (https://www.benchling.com/) and cloned into pRGE32 vector as previous described (Xie et al., 2015). Primers used for cloning are listed in Supplemental Table. *B. distachyon* transformations were performed according to *Agrobacterium tumefaciens*-mediated infection of Bd21-3 embryonic callus tissue (Vogel and Hill, 2008).

### Protein Expression and Purification

The pET151 *BdLCH1* construct was transformed into *Escherichia coli* strain BL21. A single positive clone was grown in 5 ml of LB (Luria-Bertani) medium with ampicillin (100 mg/L) overnight at 37°C. The 5 ml overnight culture was grown in 200 ml of LB medium with glucose (2 g/L) and ampicillin (100 mg/L) until OD_600_ of the culture reached 0.5. To induce expression, IPTG was added to a final concentration of 0.5 mM and cells were incubated overnight at 16°C. Cells were harvested by centrifugation and resuspended in 10 ml of binding buffer (50 mM sodium phosphate pH 8.0, 300 mM NaCl, 10 mM imidazole and protease inhibitor cocktail). Resuspended cells were then sonicated for 10 cycles of 5 seconds pulses with 15 seconds incubation on ice between pulses. The supernatant was obtained by centrifuging at 20,000g for 10 min at 4°C and then added to the HiTrap Talon crude column. After washing with 10 ml of washing buffer (50 mM sodium phosphate pH 8.0, 300 mM NaCl, 15 mM imidazole), proteins were eluted using elution buffer (50 mM Tris pH 8.0, 300 mM NaCl, 300 mM imidazole). The eluted proteins were concentrated, and the buffer was changed to 50 mM sodium phosphate pH 6.0 using Amicon ultra centrifugal filters (Sigma, Cat. No. UFC5010).

### Enzyme Analysis

The lichenase activity was measured using a commercial kit (Megazyme, K-MBG4) and the supplied protocol. 90 ul of reaction mixture was prepared after combining 30 ul of 1 mM of substrate (4,6-*O*-benzylidene-2-chloro-4-nitrophenyl-β-(3^l^-β-d-cellotriosyl-glycose) in 50% (v/v) DMSO with 60 ul of protein solution (8.4 pmol) in 100 mM of sodium acetate buffer or 100 mM of sodium phosphate buffer with various pH. The reaction was incubated in the thermocycler for 10 min at designated temperature and stopped by the addition of 2% (w/v) Tris buffer (pH 10). Lichenase activity (release of 2-chloro-4-nitrophenol) was detected at 405 nm. MBG4 unit (release of one micromole of CNP in one minute) was calculated using the equation provided with the kit.

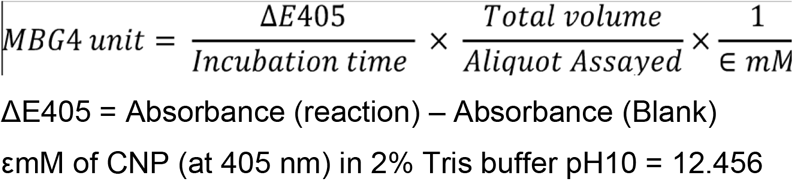

### RNAscope In Situ Hybridization

Plant samples were collected in the middle of light period and formalin-fixed in 10% NBF for 16 h at room temperature. Samples were then washed with 1X PBS and dehydrated through a series of graded ethanol washes, cleared with xylene and embedded in paraffin wax according to established protocols (Karlgren et al. 2009). Embedded *B. distachyon* samples were thinly sectioned (6 μm thick) using a Leica Microtome. Sections were placed on Fisher brand Superfrost Plus slides (Cat. No. 12-550-15) on a hot plate for 5 min and dried overnight at room temperature. Slides were processed as described (Wang et al., 2012). Specifically, RNAscope 2.5 HD Detection Reagent-RED kit was used (Cat. No. 322360). BdLCH1 specific probes were designed by Advanced Cell Diagnostics and probes that target the bacterial *DapB* gene from the kit were used as negative control. Slides were sealed with EcoMount (Cat. No. EM897L) after hybridization. Images of plant sections were obtained using bright field through a Zeiss Axio Imager M2 microscope.

### Preparation of AIR, MLG Assay, and Starch Assay

AIR was prepared as previously described (Schreiber et al., 2014) with modifications. Freeze-dried plant materials were ground into fine powder with stainless steel balls using TissueLyser (Qiagen). Ground materials were washed two times using 70% ethanol for 10 min at 97 °C followed by one wash of 100% ethanol for 10 min at 97 °C. The freeze-dried AIR was weighed, and about 3 mg of AIR was subjected to MLG or starch assays. MLG or starch was measured using a scale down version of commercially available reagents (Megazyme; MLG, K-BGLU; Starch, K-TSTA) (Mccleary and Codd, 1991; McCleary et al., 1994).

### Immunofluorescence and Immunogold labeling

Leaf or internode cross sections were prepared and immunolabeled with MLG monoclonal antibody as previously described (Fan et al., 2018; Kim et al., 2018).

### Phylogenetic Analysis

The coding sequence of putative lichenase genes from *Triticum aestivum*, *Hordeum vulgare*, *Brachypodium distachyon*, *Sorghum bicolor*, *Oryza sativa*, *and Zea mays* were obtained from Phytozome 12 (Goodstein et al., 2012; http://phytozome.jgi.doe.gov/). Evolutionary analyses were conducted in MEGA X (Kumar et al., 2018; Stecher et al., 2020). The evolutionary history was inferred by using the Maximum Likelihood method and General Time Reversible model (Nei and Kumar, 2000). The tree with the highest log likelihood (−13541.16) is shown. The percentage of trees in which the associated taxa clustered together is shown next to the branches. Initial tree(s) for the heuristic search were obtained automatically by applying Neighbor-Join and BioNJ algorithms to a matrix of pairwise distances estimated using the Maximum Composite Likelihood (MCL) approach, and then selecting the topology with superior log likelihood value. A discrete Gamma distribution was used to model evolutionary rate differences among sites (5 categories (+G, parameter = 1.0063)). The tree is drawn to scale, with branch lengths measured in the number of substitutions per site. This analysis involved 24 nucleotide sequences. Codon positions included were 1st+2nd+3rd+Noncoding. There were a total of 2623 positions in the final dataset.

### Plant Morphological Measurements

Fully senesced plants were used for morphological measurements. Plant height was measured from the base of the tallest tiller to the tip of the inflorescence. Diameter of the last-second internodes without leaf sheath was recorded as stem diameter using a digital caliper. Plant dry weight was quantified by weighing the stem and leaf tissues of senesced plants.

### Iodine Staining

Starch was visualized by Lugol solution (Sigma, Cat. No. 32922). Detached leaves from dark-incubated plants were harvested and boiled in 95% ethanol to remove pigments. After rinsing with double-distilled water, the leaves were stained with Lugol solution for 5 min. Stained leaves were rinsed with double-distilled water and then photographed using a stereo dissecting microscope with a digital camera.

### Transcriptome Profiling and Data Analysis

Total RNA preparation, library construction, multiplexed sequencing and sequence read alignment was performed as described in Fan et al. (2018).

## Accession Numbers

Sequence data for the genes mentioned in this study can be found in *B. distachyon* genome as follows: *BdLCH1* (Bradi2g27140.1), *BdCSLF6* (Bradi3g16307.1), *BdCESA1* (Bradi2g34240.1). The transcriptional profiling data used in this article can be found in the Short Read Archive at NCBI (https://www.ncbi.nlm.nih.gov/sra) and the BioSample accession numbers are as follows: EI-1, SAMN08097904; EI-2, SAMN08097907; EI-3, SAMN08097910; EI-4, SAMN08097913; EI-5, SAMN08097916; EI-6, SAMN08097919; EI-7, SAMN08097922; EI-8, SAMN08097925; EI-9, SAMN08097928; EI-10, SAMN08097931; FEI1, SAMN08097934; FEI2, SAMN08097937; FEI3, SAMN08097940; FEI4, SAMN08097943; EndoGerm, SAMN08097973; Mature leaf, SAMN08097979; Elongating leaf, SAMN10078799; Coleoptile-24h, SAMN08097868; Coleoptile-48h, SAMN08097889; Coleoptile-72h, SAMN08097898; Coleoptile-120h, SAMN08097901.

## Supplemental Data

**Supplemental Figure 1.** Protein sequence alignment of BdLCH1 with barley EI and EII.

**Supplemental Figure 2.** Activity of BdLCH1 on hydrolysis of MBG4 reagent.

**Supplemental Figure 3.** Expression profiles of *BdCESA1, BdCSLF6*, Bradi1g15295, and Bradi2g60497.

**Supplemental Figure 4.** Immunolabeling of cross sections of wild type with MLG monoclonal antibody.

**Supplemental Figure 5.** Schematic representations of targeted *BdLCH1* gene editing using CRISPR-Cas9.

**Supplemental Figure 6.** MLG accumulation in internodes of wild type and *lch1* mutants.

**Supplemental Figure 7.** Immunofluorescence and immunogold labeling of MLG in wild-type internodes.

**Supplemental Figure 8.** Plants and sampling of leaves for dark-incubation experiments.

**Supplemental Figure 9.** Expression levels of *BdCSLF6* in leaves of dark-incubated plants.

**Supplemental Figure 10.** Iodine staining of cross sections of wild-type leaves and internodes.

**Supplemental Table.** List of primers.

## ACKNOWLEDGEMENTS

We thank Nick Thrower (Michigan State University) for bioinformatics assistance in analyzing the RNA-seq data, Dr. Yinong Yang (The Pennsylvania State University) for providing plasmid pRGEB32, and Melinda Frame (Michigan State University) for help with confocal microscopy. This research was supported by the Great Lakes Bioenergy Research Center, U.S. Department of Energy, Office of Science, Office of Biological and Environmental Research under Award Number DE-SC0018409.

## AUTHOR CONTRIBUTIONS

M.F. and C.G.W. conceived the original research plans; M.F., J.K.J., J-Y.C., and C.M.B. performed most of the experiments and analyzed the data; S-J.K. and S. Z-D. performed in vitro enzyme assays and RNA in situ hybridization experiments, respectively; C.G.W. and F.B. supervised the experiments; M.F. wrote the manuscript; C.G.W., J.K.J., S-J.K., S. Z-D., and F.B. edited the manuscript.

